# A comprehensive phenotypic screening strategy to identify modulators of cargo translocation by the bacterial Type IVB secretion system

**DOI:** 10.1101/2022.02.07.479497

**Authors:** Eric Cheng, Dorjbal Dorjsuren, Stephanie Lehman, Charles L. Larson, Steven A. Titus, Hongmao Sun, Alexey Zakharov, Ganesha Rai, Robert A. Heinzen, Anton Simeonov, Matthias P. Machner

## Abstract

Bacterial type IV secretion systems (T4SSs) are macromolecular machines that translocate effector proteins across multiple membranes into infected host cells. Loss of function mutations in genes encoding protein components of the T4SS render bacteria avirulent, highlighting the attractiveness of T4SSs as drug targets. Here, we designed an automated high-throughput screening approach for the identification of compounds that interfere with the delivery of a reporter-effector fusion protein from *Legionella pneumophila* into RAW264.7 mouse macrophages. Using a fluorescence resonance energy transfer (FRET)-based detection assay in a bacteria/macrophage co-culture format, we screened a library of over 18,000 compounds and, upon vetting compound candidates in a variety of *in vitro* and cell-based secondary screens, isolated several hits that efficiently interfered with biological processes that depend on a functional T4SS, such as intracellular bacterial proliferation or lysosomal avoidance, but had no detectable effect on *L. pneumophila* growth in culture medium, conditions under which the T4SS is dispensable. Notably, the same hit compounds also attenuated, to varying degrees, effector delivery by the closely related T4SS from *Coxiella burnetii*, notably without impacting growth of this organism within synthetic media. Together, these results support the idea that interference with T4SS function is a possible therapeutic intervention strategy, and the emerging compounds provide tools to interrogate at a molecular level the regulation and dynamics of these virulence-critical translocation machines.

**Importance:** Multi-drug-resistant pathogens are an emerging threat to human health. Since conventional antibiotics target not only the pathogen but also eradicate the beneficial microbiota, they often cause additional clinical complications. Thus, there is an urgent need for the development of “smarter” therapeutics that selectively target pathogens without affecting beneficial commensals. The bacterial type IV secretion system (T4SS) is essential for the virulence of a variety of pathogens but dispensable for bacterial viability in general and can, thus, be considered a pathogen’s Achilles heel. By identifying small molecules that interfere with cargo delivery by the T4SS from two important human pathogens, *Legionella pneumophila* and *Coxiella burnetii*, our study represents the first step in our pursuit towards precision medicine by developing pathogen-selective therapeutics capable of treating the infections without causing harm to commensal bacteria.

## Introduction

To successfully establish an infection, bacterial pathogens employ a variety of strategies to evade the host immune response and to establish conditions favorable for their own survival and growth. Host cell signaling and trafficking processes are manipulated by effector proteins and microbial toxins that are encoded by pathogens and then shuttled into the host cell using specialized delivery systems. One widespread effector translocator is the type IV secretion system (T4SS) that is present in a variety of animal and plant pathogens (1, 2). Based on their architecture, T4SSs are categorized into two major classes: i) the type IVA secretion system (T4ASS) which is comprised of at least 12 different proteins and found in organisms like *Helicobacter, Brucella*, and *Bartonella;* and ii) the more complex type IVB secretion system (T4BSS), present in *Legionella pneumophila, Coxiella burnetii* and *Rickettsiella grylli*, which is assembled from twice as many components as the T4ASS (2–6). Despite their differences in composition and complexity, both classes of T4SSs are evolutionarily related to DNA conjugation systems and share a common set of subassemblies that operate in a similar manner, such as an outer membrane core complex and an inner membrane complex.

Recent advances in structural biology, electron microscopy, electron cryotomography and single-particle cryo-electron microscopy have provided much-needed insight into the three-dimensional organization of subassemblies from several T4SSs, including the VirB/VirD4 T4SS from *Agrobacterium tumefaciens* (7), the *H. pylori* Cag system (8, 9), and *Escherichia coli* conjugation apparatuses encoded by the R388 (10) and pKM101 plasmids (11–13). The Dot/Icm T4BSS system from *L. pneumophila* is one of the most-well characterized secretion system of the type IVB class (14–18). It is composed of at least ~27 components that assemble into several architectural subcomplexes, including an outer membrane core complex composed of at least five proteins (DotC, DotD, DotH, DotK, and Lpg0657) that has a pinwheel-shaped structure with a 13-fold symmetry, a periplasmic ring with an 18-fold symmetry, and an inner membrane subcomplex consisting of six proteins (DotL, DotM, DotN, IcmS, IcmW and LvgA) that traverses the inner membrane. A wide channel at the center of a stalk, composed mainly of DotG, connects the inner and outer membrane complexes of the Dot/Icm T4SS and likely ushers substrate proteins from the cytosol of the bacteria across their outer membrane. Other components function as cytosolic chaperones (IcmQ, IcmR) or as inner membrane-associated ATPases (DotL, DotB, and DotO) that convert chemical into mechanical energy for cargo translocation through the T4SS conduit. How the remaining proteins contribute to the structure and function of the *L. pneumophila* T4SS remains to be determined.

*L. pneumophila* relies on its Dot/Icm T4SS for colonization and proliferation within a wide range of host cells, including freshwater amoeba in the environment and alveolar macrophages during Legionnaires’ pneumonia in humans (19). Throughout its intracellular replication cycle, the bacterium resides within a membrane-enclosed compartment, the *Legionella*-containing vacuole (LCV), that avoids fusion with destructive endosomes and lysosomes (20, 21). Instead, the bacterium acquires proteins and membranes from the early secretory pathway and other sources to establish a camouflaged replication compartment (22). *L. pneumophila* mutants with a non-functional T4SS fail to control trafficking of their LCV and are quickly delivered to lysosomes for degradation (23–25), thus underscoring the importance of the Dot/Icm system for *Legionella* pathogenesis.

Given their importance for virulence, bacterial type IV secretion systems are being increasingly recognized as putative drug targets. Since T4SSs are typically not essential for bacterial fitness outside the host, they experience low selective pressure in the environment, making it less likely for resistance mechanisms against inhibitory compounds to already exist or to easily spread throughout microbial populations. Moreover, since commensal bacteria within the microbiota of humans are not known to require a T4SS for survival, therapeutics that specifically target these translocation machines will likely affect only pathogens while leaving the healthy microbiota undisturbed, thus reducing the risk of secondary infections by creating niches for pathogens.

Recently, several groups have reported the discovery of small molecules that interfere with the activity of type III secretion systems (T3SSs) (26), another major bacterial translocation machine found in pathogens. While the exact type of reporter assays used in these studies varied, they all made use of the convenient fact that T3SS activity could be monitored during bacterial growth in broth without the need for host cells to be present. This is in stark contrast to bacterial T4SS that do not exhibit effector translocation activity during axenic growth and that are activated only upon contact of the pathogen with a target cell, which can either be another bacterium in the case of T4SS-mediated DNA conjugation, or a host cell during infection (27). The requirement for both bacteria and host cells to be present at the same time and to interact in a productive manner that allows for efficient reporter transfer to occur explains why high throughput screens for compounds that interfere with T4SS function have remained a major challenge.

To bypass these limitations, alternative approaches have been developed, for example by identifying inhibitors of individual protein components of T4SSs, such as *Brucella* VirB8 (28, 29) or *H. pylori* VirB11 (30). While these surrogate screening approaches have made some progress towards the discovery of T4SS inhibitors, they did not fully recapitulate the complex behavior of host-pathogen interactions during infection. Shuman and colleagues (31) have made a first progress towards identifying molecules that attenuate the delivery of a effectors from *L. pneumophila* into infected host cells. In their screening campaign, they tested a library of approximately 2,600 compounds and identified several candidates that reduced delivery of a reporter-effector fusion protein by *L. pneumophila* into J774 macrophages. Importantly, the vast majority of hit compounds that were detected in this way appeared to indirectly interfere with processes on the host side, most notably actin polymerization which is a dynamic process critically important for bacterial uptake via phagocytosis. By blocking phagocytosis, these compounds likely prevented the establishment of an intimate cell-cell contact between the bacterium and its host, thus reducing the efficiency of reporter protein translocation. Although valuable as tools for the experimental analysis of T4SS function, compounds that target human cell are undesirable as therapeutics against infectious diseases due the likelihood of them causing side effects.

Here, we implemented a modified screening approach to identify novel compounds that interfere with the process of reporter translocation without affecting phagocytosis. After screening a library of more than 18,000 compounds at a wide range of concentrations, several hits emerged that showed high efficacies in a variety of biological assays that require a functional T4SS, indicating that our approach was successful in selecting potentially novel therapeutics to antagonize bacterial infections by *L. pneumophila* and, notably, also *Coxiella burnetii* whose virulence relies on a related T4BSS.

## Results

### Development of a cell-based reporter assay for monitoring bacterial effector translocation

To identify compounds that specifically interfere with the process of bacterial type IV secretion, we developed a screening protocol that minimized the enrichment of compounds that function by altering host cell physiology and that favored detecting compounds that block other aspects of effector translocation by the T4SS. For this purpose, several important criteria and screening parameters had to be tested and optimized, including: the type of reporter enzyme and the substrate used for its detection; the type of host cell that, upon contact with *L. pneumophila*, favors reporter translocation under infection conditions; the ratio of bacteria to host cells and the duration for which they will be incubated; and the concentration and incubation periods of compounds from the screening libraries.

Prior to our study, several reporter protein-based translocation monitoring assays have been successfully used in *L. pneumophila*. These included the reporter enzyme Cre recombinase from P1 bacteriophage (32), the adenylate cyclase CyaA from *B. pertussis* (33, 34), and the *Enterobacteriaceae* β-lactamase (βLac) (35). The CyaA and Cre recombinase assays require either extensive washing steps or a time-intensive reporter step which makes them less desirable for a high-throughput screen (36). The βLac translocation assay bypasses many of these limitations since no wash steps are required, the experimental procedure can be completed in a single day, and the time between the addition of the reporter substrate and the readout occurs within a few hours. When fused to a known translocated substrate of the Dot/Icm T4SS, the βLac reporter is shuttled into target cells where its presence can be optically detected using CCF4/AM, a membrane-permeable lipophilic ester that is easily taken up by cultured cells (Figure 1A) (37). CCF4/AM is composed of two fluorophores (7-hydroxycoumarin and fluorescein) that form a fluorescence resonance energy transfer (FRET) pair linked by a β-lactam ring. The excitation of intact CCF4/AM with light of 409 nm wavelength results in FRET from the coumarin donor to the fluorescein acceptor, causing cells without βLac to emit light in the green spectrum (518 nm). In the presence of βLac activity, CCF4/AM cleavage occurs which separates the two fluorophores and disrupts FRET, resulting in cells that fluoresce in the blue spectrum (447 nm) of the coumarin donor (Fig. 1A). The ratio of blue-to-green light in infected cells is an approximation of the level of translocated effector-reporter fusion protein and can serve as an indicator for the activity of the *L. pneumophila* T4SS.

**Figure 1:**
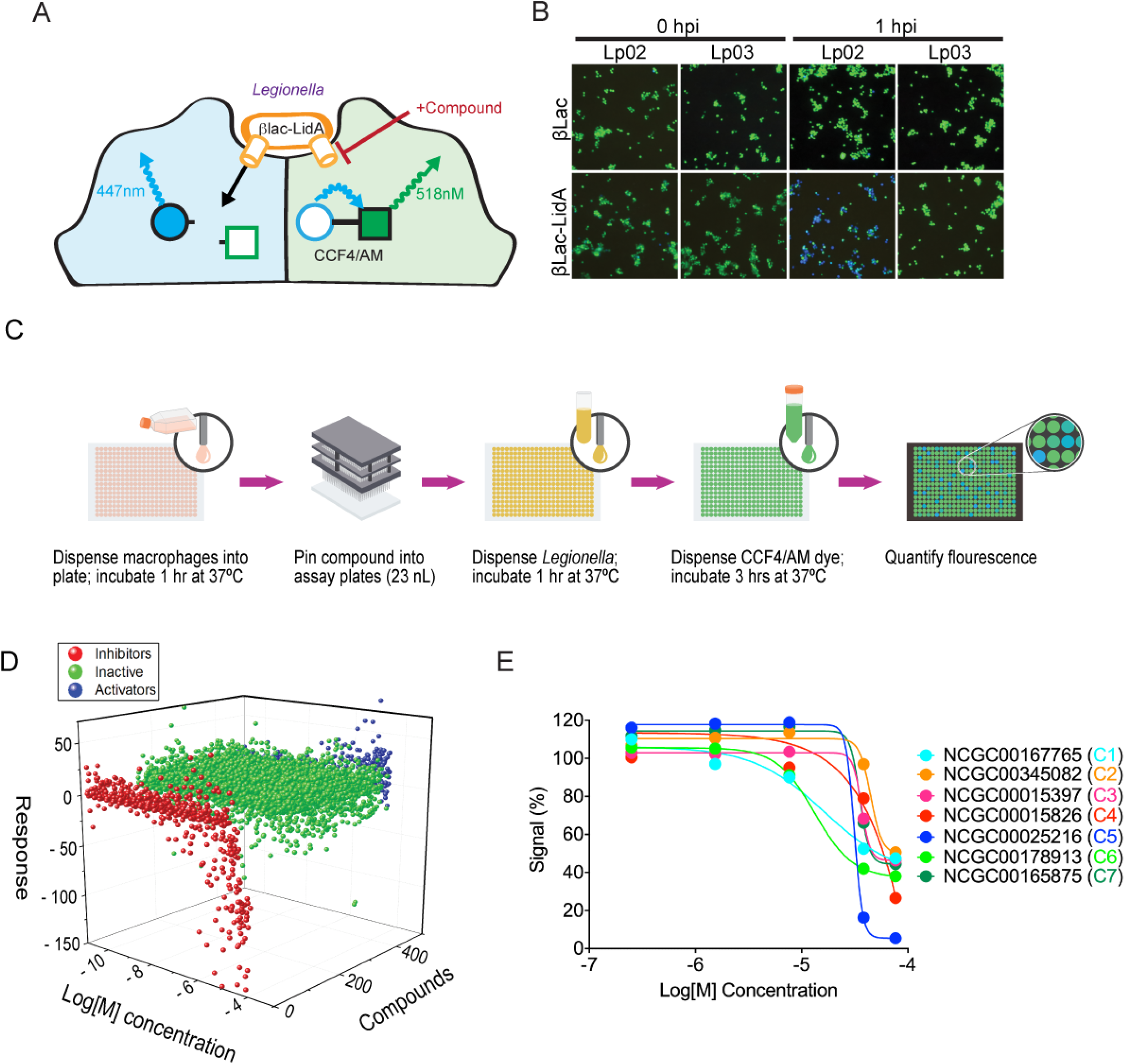
Small molecule library screen using a FRET-based reporter assay. (A) Schematic depiction of the reporter translocation assay. Compounds that interfere with T4SS-mediated reporter delivery into infected cells will prevent β-Lac-mediated cleavage of the mammalian cell-permeable CCF4/AM substrate, retaining its green fluorescence, whereas reporter delivery into host cells will result in a shift in emission light from green to blue due to cleavage of the FRET pair. (B) RAW264.7 macrophages were challenged at an MOI of 20 with either Lp02 (functional T4SS) or Lp03 (defective T4SS) harboring plasmids encoding either βLac (control) or βLac-LidA. At the indicated time points (hours post infection (hpi)), CCF4/AM was added to the cells, and fluorescence emission light was detected by epifluorescence microscopy. (C) Schematic overview of the different high-throughput screen stages. (D) Waterfall plot depicting the dose response to all compounds. Compounds that result in reduced reporter translocation (assessed by FRET) are shown in red, compounds without effect are shown in green, activators or false activators/fluorescent compounds are shown in blue. E) Inhibition profile for representative hits from the FRET confirmation assay using an 11-point dose range. Data were globally fit using a Hill equation.

We created a genetic fusion between βLac and LidA, a known substrate of the *L. pneumophila* T4SS and a protein shown to efficiently shuttle βLac into host cells (38). Synthesis of equal levels of βLac-LidA in both *L. pneumophila* wild-type (Lp02) and the T4SS-defective *L. pneumophila dotA3* (Lp03), a strain with a nonfunctional variant of the polytopic inner membrane protein DotA (39, 40), was confirmed by immunoblot analysis (Supplemental Fig. 1). Using the FRET-based read-out, we detected efficient translocation of βLac-LidA during infection of mouse RAW264.7 macrophages, a cell type that can be easily propagated and readily takes up *L. pneumophila* (Fig. 1B) (36). Infection with the Lp02 strain producing βLac-LidA resulted in efficient hydrolysis of the CCF4/AM substrate within RAW264.7 macrophages, causing them to fluoresce blue, whereas no hydrolysis was detected upon macrophage challenge with either Lp03 producing βLac-LidA or Lp02 producing unconjugated βLac (Fig. 1B). Thus, reporter translocation depended on its fusion to a translocated effector and the presence of a functional Dot/Icm system.

One of the critical aspects of the high-throughput library screen was the miniaturization of the FRET assay from a 96-well to a 1536-well plate format. This increases the number of samples that can simultaneously be screened in each plate by 16-fold and, at the same time, reduces the total sample volume per well from 220 μl to 6 μl. Despite these benefits, miniaturization also is associated with a variety of challenges, most notably enhanced fluid surface tension, limited sample mixing or liquid aspiration capability, as well as liquid evaporation and an increased surface-to-volume ratio that can dramatically affect reagent absorption and stability (41). To assure assay reproducibility under these difficult conditions, the translocation assay described above (Fig. 1A) was further optimized for a 1536-well plate format. We found that a bacteria-to-macrophage ratio of 20:1 and an infection period of 60 minutes produced the best results, with an average Z’ factor (42) of 0.63±0.14, which exceeded the generally accepted criteria of a Z’ factor >0.5 required for high-throughput screening. These results suggest a reproducible assay that is not adversely affected by minor variabilities in experimental conditions.

### High-throughput screen (HTS) for compounds that attenuate reporter delivery by the *L. pneumophila* T4SS

Once optimized for a 1536-well plate format, the FRET-based translocation assay was carried out by performing qHTS against commercially available small libraries (see Materials and Methods) at multiple concentrations, to evaluate compounds for their ability to interfere with T4SS-mediated β-Lac-LidA translocation (Fig. 1C). A total of 18,272 compounds were initially screened at either five or six concentrations ranging from 22 nM to 46 μM (43–46). Importantly, compounds and bacteria were added to macrophage monolayers in very short succession (<15 minutes) as to minimize effects of the compounds on host cell physiology. A dose-response curve was generated for each compound and classified to one of four curve classes based on the shape of the curve and the quality of the fit (*r*^2^) (45) (Fig. 1D,E). Briefly, curve class 1 represents a full curve with full efficacy range and high degree of fit (high R^2^), curve class 2 represents curve with partial efficacy and lower degree of fit (low R^2^), class 3 is a dose response with a single-point trend, while class 4 is a flat, or inactive, response. We represented the screening outcome in three categories: all actives shown in red were compounds that inhibited in the assay (negative curve classes 1-3), inactives were in green (class 4), and in blue we displayed compounds that appeared to activate in the assay (positive curve classes) (Fig. 1D). Of the 18,272 compounds evaluated in the primary screen, 501 compounds were active with a maximum response (≥35% signal reduction), corresponding to a hit rate of 2.7 % (Fig. 2).

**Figure 2:**
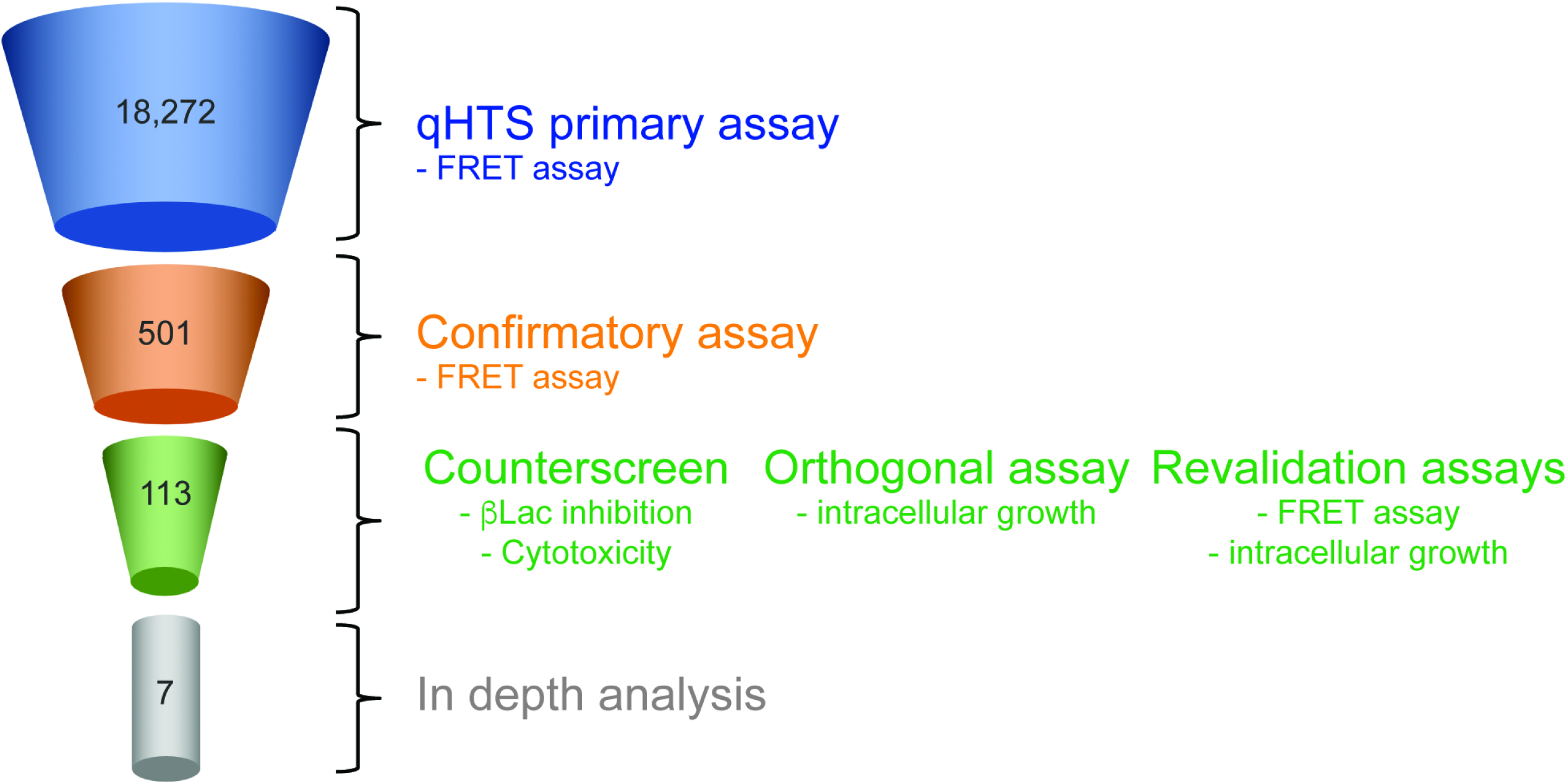
Compound triage. Schematic overview of the multi-step compound triage workflow. Numbers represent the quantity of compounds that emerged from each triage step.

Active primary hits were further evaluated in the same primary assay using an 11-point dose response, and 113 compounds (0.6%; Table S1) were confirmed as positive hits (Fig. 2). These compounds were further assessed in counter-screens and orthogonal assays as described next.

### Triaging of active compounds

Given the complexity of the bacteria-to-macrophage translocation process, several irrelevant mechanisms could have caused a reduction in the FRET-based reporter signal without directly affecting T4SS-mediated translocation. These include a potentially inhibitory effect of hit compounds on the enzymatic activity of β-Lac; compounds that cause host cell toxicity or lysis; and compounds that are highly promiscuous and thus non-specifically react with macromolecules. Each of these issues was addressed using the following validation methods.

First, we determined if any of the active compounds functioned as β-Lac inhibitors that interfered with the enzyme’s ability to hydrolyze its substrate. To do so, we adapted a chromogenic assay that monitors the ability of β-Lac to hydrolyze the cephalosporin-type substrate nitrocefin in solution (47) (Supplemental Fig. 2A). Upon hydrolysis of the amide bond in its β-lactam ring, nitrocefin undergoes a rapid and distinctive color change from yellow (390 nm) to orange-red (486 nm) that can easily be measured using a spectrophotometer. Sixteen of the 113 compounds blocked β-Lac enzymatic activity, of which three were known inhibitors of β-Lac (Pivoxil Sulbactam (NCGC00249610), Tazobactam sodium (NCGC00159340), and Clavulanate lithium (NCGC00180892)).

We also explored the possibility if any of the active compounds affected mammalian cell viability, which could have prevented uptake and/or conversion of CCF4/AM during the FRET-based assay. Moreover, cytotoxicity is an undesirable trait in therapeutics as it increases the chance of side effects. To determine cell viability, RAW264.7 macrophage monolayers were incubated for 24 hours with hit compounds at concentrations ranging from 22 nM to 46 μM. The cells were subsequently lysed, and ATP levels, a proxy for live cell metabolism, were measured using a luminescence-based signal that is proportional to the amount of ATP present (Supplemental Fig. 2B). Of the 113 active compounds selected from the primary assay (Table. S1), 62 compounds had only a moderate or no effect on cell viability (all negative curve classes and max response more than 35) even after 24 hours of incubation and were advanced to the next stage. To eliminate non-specific inhibitors, we deployed an orthogonal GFP intracellular growth *L. pneumophila* assay in macrophages as described below.

### Compound treatment protects macrophages from intracellular growth of *L. pneumophila*

Given that effector translocation is essential for *L. pneumophila* intracellular survival and growth, we speculated that treatment of *L. pneumophila* with hit compounds that affect T4SS function should render the bacteria less virulent and protect host cells from intracellular bacterial replication (Fig. 3A). We developed a 1536-well plate-based automated microscopy and image processing approach to monitor growth of GFP-producing *L. pneumophila* within host cells. After pre-treatment with compounds (22 nM to 46 μM final concentration) for 15-30 minutes, RAW264.7 macrophages were challenged with a GFP-producing *L. pneumophila* strain. After incubation for 14 hours, which is equivalent to the time required by *L. pneumophila* to complete its intracellular replication cycle, macrophages were chemically fixed, host cell nuclei were labeled by staining DNA with the dye Hoechst-33342, and cells were imaged using an INCell Analyzer 2200 Imaging system to identify cell nuclei and GFP-positive bacteria (Fig. 3B,C). INCell analyzer software (GE Healthcare Life Sciences) was used to quantify cells with nuclei stain. GFP intensity was normalized to DMSO control to identify the cells with positive-GFP. Of 62 compounds tested in our *L. pneumophila* intracellular growth assay in conjunction with the β-Lac assay resulted in 33 compounds that demonstrated inhibition with an efficacy greater than 60% and AC_50_s lower than 25 μM (Supplemental Fig. 3). Of these, we manually excluded 11 compounds with known antibacterial activities and mercury containing promiscuous compounds. The remaining 22 compounds were re-acquired from a commercial source, purified, and re-tested in 384-well format to validate previous results. Of these 22 compounds, seven compounds (NCGC-00167765, −00345082, −00015397, −00015826, −00025216, −00178913, −00165875, called C1 to C7 hereafter) were promoted for further evaluation (Fig. 3D; Table S2).

**Figure 3:**
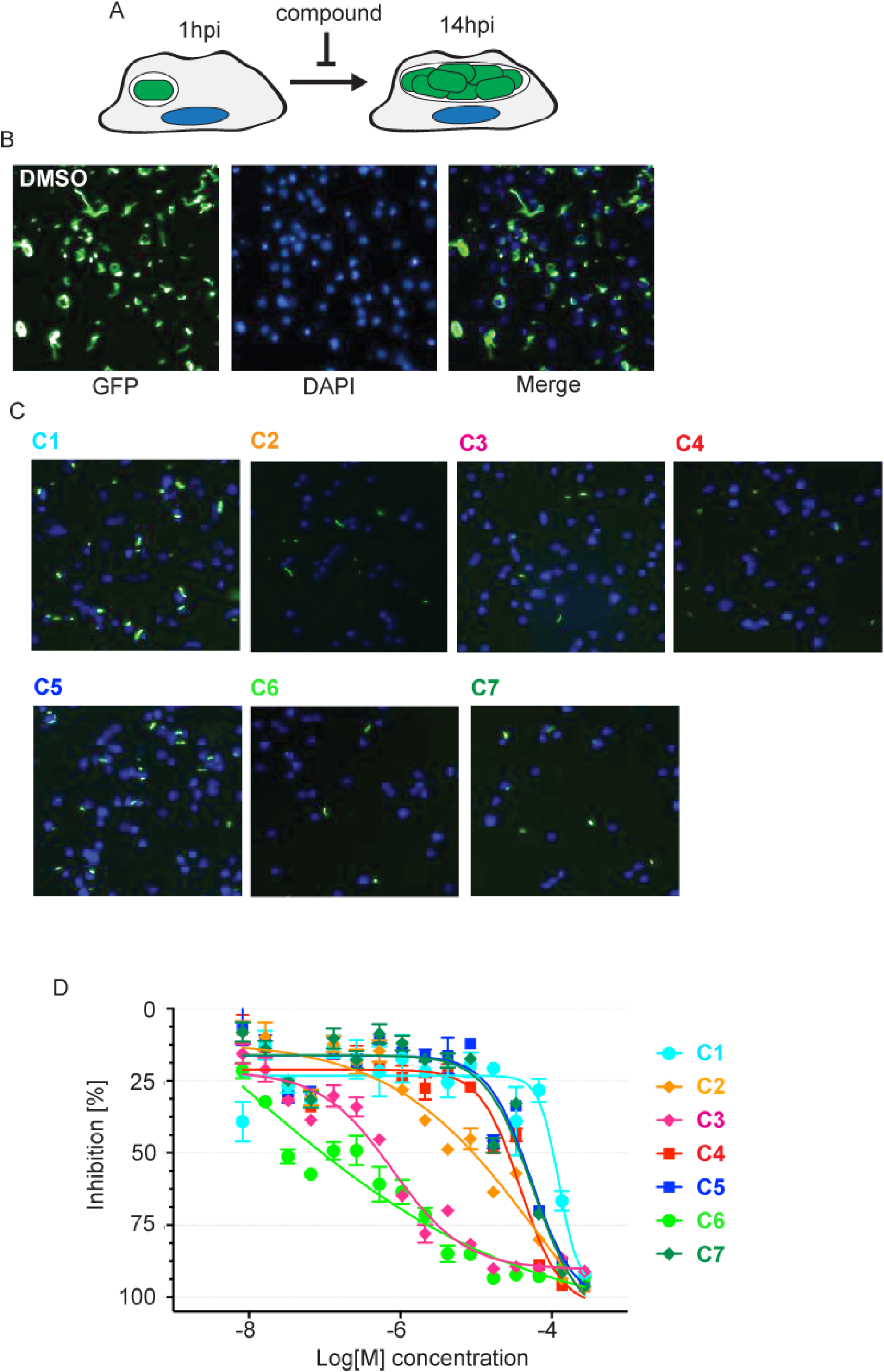
*L. pneumophila* intracellular growth is attenuated in the presence of hit compounds. (A) Schematic overview of the intracellular growth assay. (B) Growth of *Lp02ΔflaA* in RAW264.7 macrophages. RAW264.7 macrophages were challenged with Lp02 in the presence of DMSO (vehicle). Images were captured using an InCell Analyzer at 14 hours post infection (hpi). Bacteria are shown in green and DNA (DAPI staining) in blue. (C) Growth of Lp02 in RAW264.7 macrophages is attenuated in the presence of compounds. RAW264.7 macrophages were challenged as described in (A) in the presence of the indicated compounds at 57 μM (NCGC IDs shown). Panels are merged images of GFP-producing *L. pneumophila* (green) and DAPI-stained host nuclei (blue). (D) Inhibition of *L. pneumophila* growth by select compounds is dose-dependent. In RAW264.7 macrophages, *L. pneumophila* growth was quantified by measuring the GFP signal strength relative to macrophages not infected by *GFP*-Lp02. Experiments were done in triplicate.

### Studying the effect of hit compounds on bacterial uptake

Efficient effector translocation requires close contact between bacteria and their host cell. This condition is strongly favored during phagocytosis where the pathogen is being engulfed by the host plasma membrane. Many of the compounds previously shown to block T4SS-mediated effector translocation indirectly did so by attenuating *L. pneumophila* uptake (31). Although the screening approach implemented here had been designed to reduce the enrichment of compounds that interfere with host cell processes, we nonetheless evaluated if and to what extent the hit compounds negatively impacted phagocytosis of *L. pneumophila.* RAW264.7 macrophages were challenged for 1 hour with Lp02 in the presence of either the vehicle (DMSO), hit compounds C1 to C7, or cytochalasin D, a cell-permeable and potent inhibitor of actin polymerization, and the percentile of cells with intracellular bacteria was microscopically determined (Supplemental Fig. 4). Unlike cytochalasin D, which reduced the number of intracellular bacteria by almost 95%, four of the seven hit compounds had no significant effect on *L. pneumophila* phagocytosis. Compound C4 and C5 still allowed 59% and 71% of bacterial uptake by macrophages, respectively, while C2 more strongly attenuated phagocytosis by -65%. Thus, of the seven hit compounds tested here, six showed moderate to low effects on bacterial phagocytosis, confirming that our screening protocol had indeed favored the identification of hit compounds that attenuate effector delivery into host cells by some other means, a notable difference to the aforementioned study by Shuman and colleagues (31).

### Effect of hit compounds on *Legionella* growth outside of host cells

The inhibitory effect of the seven selected hits on *L. pneumophila* growth in RAW264.7 macrophages might suggest that they could have attenuated effector protein translocation through the T4SS, its most important known function in host intracellular growth. Alternatively, these compounds could also exhibit bacteriostatic or bactericidal activity, comparable to the role of conventional antibiotics, which would reduce the ability of *L. pneumophila* to proliferate within host cells. To distinguish between these two possibilities, we measured the effect of the seven hit compounds on growth of Lp02 outside its host, conditions under which the T4SS can be considered dispensable. The bacteria were incubated in AYET media containing either DMSO (vehicle), the antibiotic chloramphenicol (5 μg/ml), or C1 to C7 (28μM final concentration), and growth was monitored by measuring the optical density at 600 nm (OD_600_) (Fig. 4A). Over a period of 18 hours, Lp02 grew robustly and uninhibited in the presence of the DMSO vehicle, whereas no increase in cell numbers was detectable in AYET media supplemented with chloramphenicol. Five compounds had no detectable effect on growth of Lp02 in AYET, suggesting that they did not have a bacteriostatic or bactericidal effect on *L. pneumophila* (Fig. 4A). Notably, two compounds C2 and C6 did cause a dramatic reduction in growth similar to the level of inhibition observed upon cultivation of the bacteria with chloramphenicol. Unexpectedly, we found that, under similar growth conditions, neither compound had an inhibitory effect on the replication of two other gram-negative bacteria, *E. coli* and *Pseudomonas aeruginosa* (Fig. 4C, D), suggesting that these two compounds did not function as general gram-negative antibiotics but were directed primarily against processes or components present in *L. pneumophila* while absent from *E. coli* and *P. aeruginosa*.

**Figure 4:**
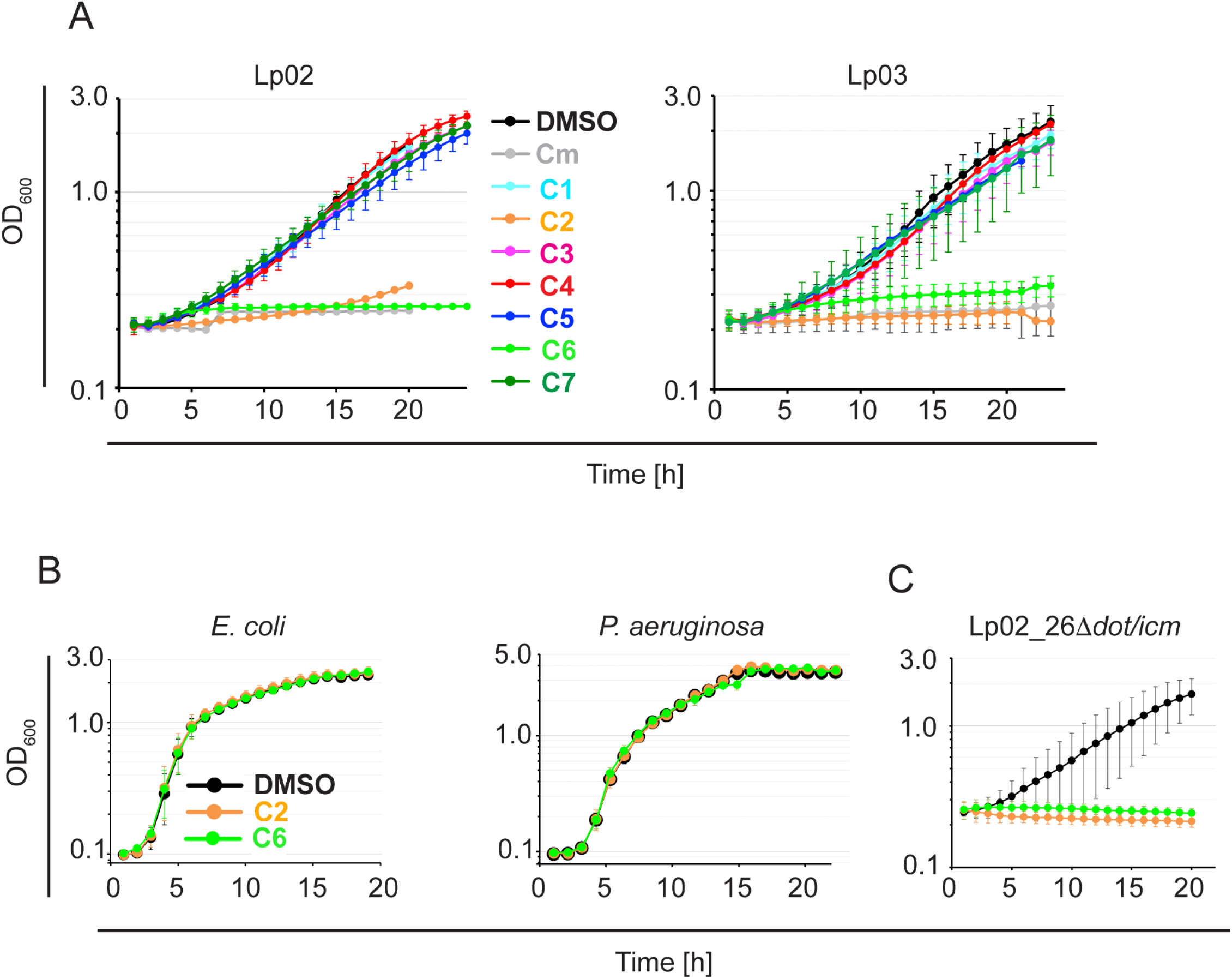
Effect of hit compounds on growth of *L. pneumophila* in liquid media. (A) Growth of *L. pneumophila* Lp02 or Lp03 in the presence of hit compounds. Bacteria were inoculated in AYET media at an OD_600_ of 0.1 in the presence of either DMSO, the antibiotic chloramphenicol (control; 5ug/ml), or the indicated compounds (NCGC IDs 28μM). Growth was monitored for at least 18 hours by measuring the absorbance at OD_600_. (B) Compounds C2 and C6 do not affect growth of *E. coli* MG1655 or *Pseudomonas aeruginosa*. (C) C2 and C6 do inhibit growth of *Lp02_26Δdot/icm*.

Although it is generally accepted that the *L. pneumophila* Dot/Icm T4SS transport system is dispensable for axenic reproduction in growth media, there are reports where a dysregulated T4SS system can affect *L. pneumophila* physiology and proliferation outside its host. Specifically, a functional T4SS renders *L. pneumophila* sensitive to high concentrations of sodium chloride within the growth media, while isolates with a non-functional T4SS, such as the Lp03, are salt-resistant (48). Although the molecular details underlying this Dot/Icm-dependent salt-sensitivity in growth media have yet to be determined, it is likely that the intact T4SS translocation pore of *Legionella* may allow unregulated passage of ions across the bacterial membrane, thus disturbing the bacterium’s electrochemical gradients. To determine if C2 and C6 had caused a similar disturbance by targeting the integrity of the Dot/Icm system, we tested whether growth in media could be restored by disabling the T4 translocon. Interestingly, we found no difference in sensitivity between Lp02 and Lp03 to either C2 or C6 during growth in AYET media (Fig. 4B). In fact, even the axenic growth of a *L. pneumophila* strain lacking the entire *dot/icm* gene cluster (Lp02(Δ26)) was robustly repressed in the presence of either C2 or C6 but not of the vehicle DMSO (Fig. 4E), demonstrating that the growth-inhibitory effects of these two compounds were independent of components of the Dot/Icm system.

### Treatment with hit compounds increases delivery of *Legionella* to lysosomal compartments

Failure to translocate effectors results in *L. pneumophila* to be rapidly shuttled to lysosomal compartments for degradation. To evaluate the effect of the five remaining hit compounds (C1, C3, C4, C5, C7) on intracellular trafficking, RAW264.7 cells were challenged with *L. pneumophila* producing mCherry, and LCVs were enumerated by indirect immunofluorescence microscopy for their colocalization with lysosomal marker protein Lamp1 (Fig 5). While *L. pneumophila* showed the commonly observed basal level (~15%) of Lamp1-positive LCVs in vehicle-treated cells, five of the six hit compounds caused a two-fold or higher increase in the number of Lamp1-positive LCVs, with C4 having the most dramatic effect with up to 80% Lamp1-positive LCVs. Combined with the results from the intracellular growth analysis and the reporter translocation assay, these data favor the scenario where the hit compounds attenuate *L. pneumophila* intracellular growth and lysosomal avoidance by altering the dynamics of effector translocation through the T4SS.

**Figure 5:**
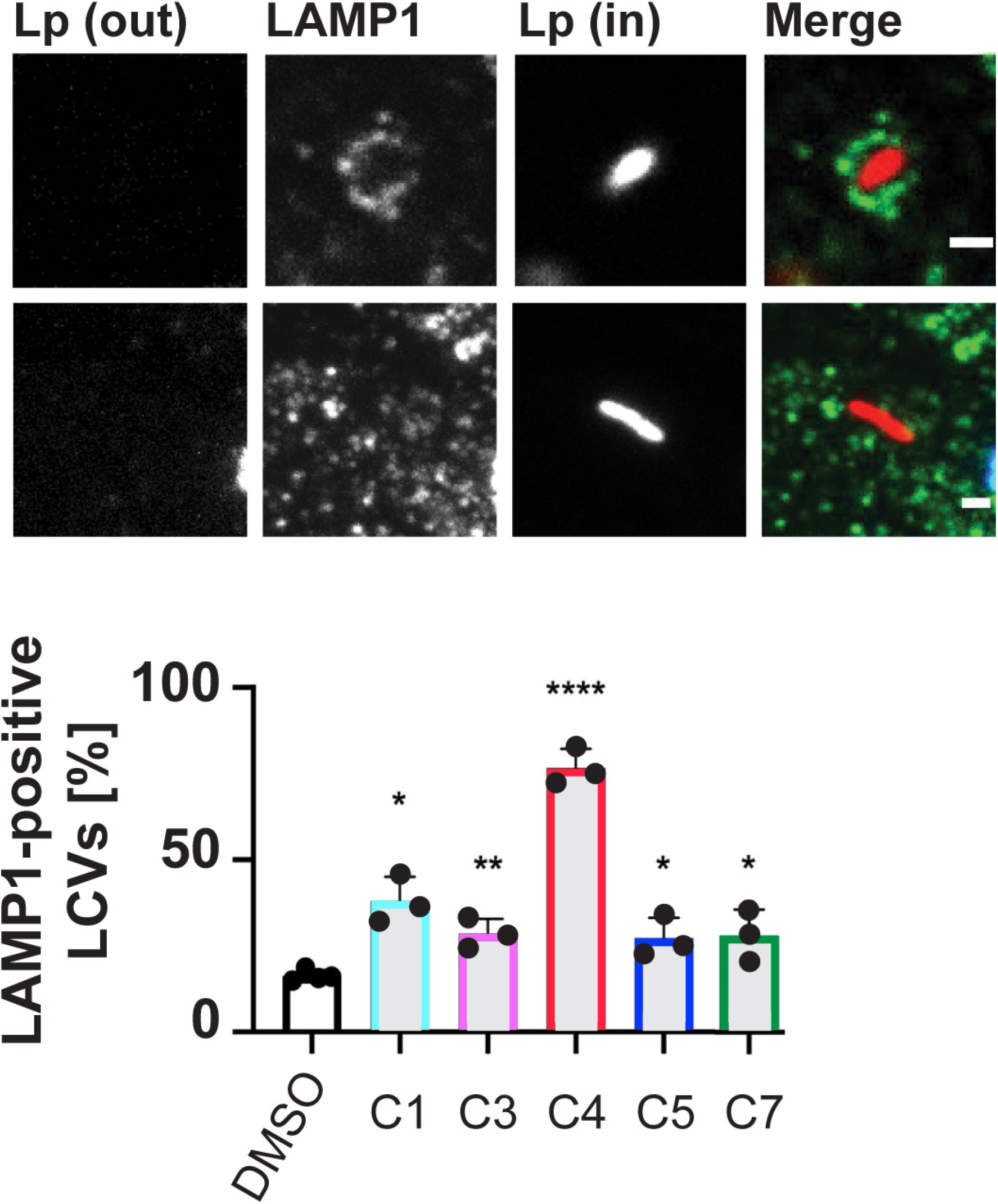
Increased delivery of *L. pneumophila* to Lamp1 positive compartments in the presence of compounds. RAW264.7 macrophages were infected with Lp02-mCherry and indicated compounds or DMSO vehicle for 2 hpi at an MOI of 25, then enumerated for the percent of internalized bacteria in Lamp1 positive compartments. (A) Representative image of Lamp1 (green) positive internalized *L. pneumophila* (red), and (B) Lamp1 negative internalized *L. pneumophila*. (C) Enumeration of Lamp1 positive *Legionella* containing vacuoles (LCVs). Data are represented as the mean of 3-4 experimental replicates with standard deviation and individual replicate points shown. (*) indicates P<0.05 compared to DMSO control.

### Hit compounds interfere with the delivery of Coxiella T4SS effector-reporter fusions

*Coxiella burnetii* is the causative agent of zoonotic Q fever in humans. Upon internalization by a permissive macrophage, the obligate intracellular bacterium resides within the *Coxiella*-containing vacuole (CCV). *C. burnetii* encodes a T4SS that is homologous to the Dot/Icm system from *L. pneumophila* that delivers bacterial effector proteins across the CCV membrane into the host cytosol (4, 49, 50). Here, we tested whether compounds that interfere with the *L. pneumophila* T4SS also inhibit translocation of *Coxiella* Dot/Icm effectors CvpA and CvpB. THP-1 macrophages were cultured for 24 h with *C. burnetii* expressing the Dot/Icm effector CvpA or CvpB fused to the CyaA reporter tag and then incubated an additional 24 h with 50 μM of each of the five indicated compounds. Cytosolic cAMP measured within infected cell lysates was used to determine the translocation efficiency of the CyaA fusions in response to compound treatment. With the exception of C4 each of the compounds reduced the cAMP levels at least 10-fold compared to cells without inhibitor treatment, both for cells infected with *C. burnetii* expressing either CyaA-CvpA or CyaA-CvpB (Fig. 6). In all instances, low levels of translocated protein were still detected above the 2.5-fold cutoff for background signal. This response could be due to persistent low levels of secretion in the presence of inhibitors or active remnants of CyaA fusions that were delivered to the host cell cytosol prior to addition of the inhibitor compounds. Importantly, none of the compounds had an inhibitory effect on *C. burnetii* growth in axenic media (Supplemental Fig. 5), further supporting the idea that they did not cause a global disturbance within the bacteria’s physiology. Together, these result support the conclusion that the inhibitor screen performed here succeeded in identifying compounds that likely interfere with cargo translocation by the bacterial T4BSS found in pathogens like *L. pneumophila* and *C. burnetii.*

**Figure 6:**
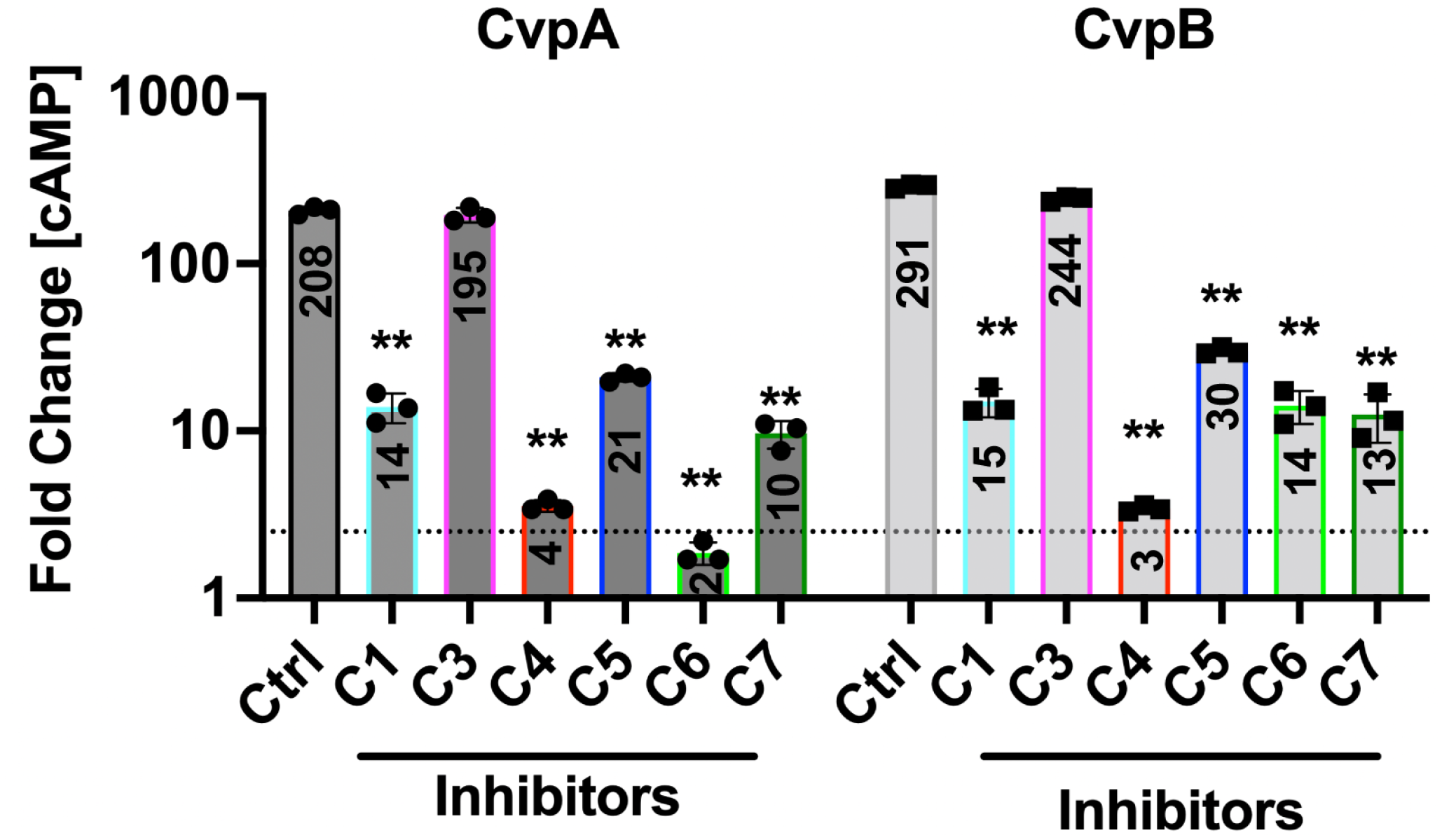
Compound treatment attenuates effector translocation by *C. burnetii* Dot/Icm. Histograms depict the fold change in cytosolic [cAMP] for THP-l cells infected for 48 hours with *C. burnetii* producing CyaA-CvpA or CyaA-CvpB fusion proteins. The cutoff for positive secretion is indicated by a dotted line at 2.5-fold change [cAMP]. Results are representative of three independent experiments and error bars indicate the standard deviations from triplicate samples. Asterisks indicates a statistically significant difference (P < 0.01).

## Discussion

In this study, we have identified and experimentally validated five hit compounds that interfered with processes that depend on a functional T4BSS, such as the delivery of a β-lactamase reporter into host cells (Fig. 1A), the proficiency of *L. pneumophila* to avoid endolysosomal trafficking (Fig. 5), and the capability to replicate within human macrophages (Fig. 3A). Importantly, none of the five compounds affected the ability of either *L. pneumophila* (Fig. 4A) or *C. burnetii* (Supplemental Fig. 5) to replicate outside their host when grown within synthetic media, conditions under which a functional Dot/Icm system is dispensable. Together, these results suggest that the five hit compounds, by targeting either the bacterial T4SS or factors required for its activation or function, attenuate *L. pneumophila’s* ability to fully employ this translocation system to deliver cargo into recipient cells.

The T4SS inhibitors described here emerged from a high throughput small molecule library screen in which we employed a protocol that strongly favored the discovery of compounds that likely function by directly altering the regulation and/or function of the Dot/Icm system instead of compounds that indirectly affect reporter delivery, for example, by altering host cell processes such as phagocytosis, a process known to be essential for *L. pneumophila* uptake. This was accomplished by adding both the bacteria and each compound in rapid succession to macrophage monolayers, thus limiting the exposure time of cultured macrophages to them. Another feature of our screening protocol was that each compound was tested at a wide range of concentrations, which allowed us to generate a dose response profile for each compound (Fig. 1D,E). As a result, our screen yielded a diverse set of hits and was enriched in compounds that seemed to affect T4SS function without altering host cells processes like phagocytosis (Supplemental Fig. 4) or cell viability in general (Supplemental Fig. 2B).

The structures of the hit compounds and their pharmacological properties are summarized in Table S2. Briefly, among these five hits, C7 (PAP-1, muscarinic acetylcholine receptor M1 antagonist), C5 (roxindole, serotonin 1a receptor agonist), C1 (LP-44, sigma1/2 receptor agonist) and C3 (ethamivan, 5-HT7 receptor agonist) are associated with modulation of listed G protein-coupled receptor (GPCR) activities, while C4 (perphenazine) is a known disruptor of HIF-β/ Transforming Acidic Coiled Coil Containing Protein 3 (TACC3) complex in the HIF pathway. Importantly, while each of these compounds has a known host cell target, we do not imply that engagement of those targets was responsible for the inhibitory effect on cargo translocation by the *L. pneumophila* T4SS. Instead, these compounds may have additional targets on either the bacterial side or the host cell side that were responsible for inhibiting effector translocation, and it will be interesting to determine the identity of those alternate targets in the future.

The fact that the hit compounds identified here show a wide range of molecular structures and chemical properties suggests that they may have different modes of inhibition, perhaps offering the opportunity to explore future dual synergistic anti-microbial actions. This is further supported by the variation in lysosomal delivery of *L. pneumophila* upon compound treatment, where the bacteria were trafficked with higher frequency to destructive lysososmal compartments within infected host cells (Fig. 5). One obvious possibility would be for hit compounds to sterically obstruct the T4SS channel, thereby preventing cargo from entering or passing through the translocon. A compound might also block a yet-to-be identified sensor on the bacterial surface or its cognate host cell receptor, thus preventing their engagement which would otherwise trigger T4SS activation upon bacterial contact with and engulfment by recipient cells. Membrane-permeable compounds might enter the bacterial cell and target cytosolic or integral membrane components of the Dot/Icm system, including subunits with ATPase activity, such as DotL or DotB (51, 52), which would disable the force-providing components of the T4SS. And, some of the compounds could bind to and block the function of the coupling protein complex composed of DotL, DotM, and DotN, which serve as a platform for the recruitment of effector proteins prior to translocation (53). Despite the briefness of the exposure of the bacteria to compounds prior to bacterial contact with macrophages, it cannot be excluded that the compounds functioned by inhibiting expression of *dot/icm* genes or by preventing proper assembly of the pre-synthesized T4SS components. This is particularly true during *Coxiella* infection where incubation periods with the compounds were prolonged compared to the *Legionella* infection assay and where effects of the compounds on the assembly of the T4SS cannot be excluded (Fig. 6).

Hit compound C4 emerged as a particularly promising candidate for interfering with T4SS-based virulence processes of *Legionella*. This compound robustly decreased the ability of *L. pneumophila* to escape lysosomal compartments (Fig. 5) and to proliferate within mouse macrophages (Fig. 3). It also dramatically reduced reporter-effector translocation by the *L. pneumophila* Dot/Icm system (Fig. 1E). Importantly, while C4 had no detectable effect on effector-reporter delivery by *C. burnetii* (Fig. 6), the four other hit compounds, C1, C3, C5, and C7, did, reducing cAMP levels by at least 10-fold compared to untreated cells (Fig. 6). Despite the high degree of homology between the T4SS components from *L. pneumophila* and *C. burnetii*, the two pathogens show notably different infection dynamics: While *L. pneumophila* effector translocation begins immediately upon host cell contact, *Coxiella* are metabolically inactive at early times post infection and do not secrete proteins until 24-48 hours later. Consequently, the different treatment regimens that were used may explain the observed difference in compound efficacies; while compounds were simultaneously added with *L. pneumophila* bacteria to host cell monolayers, macrophages infected with *Coxiella* were not exposed to the compounds until 24 hours after the initial bacterial challenge.

We also discovered in our screen two compounds C2 and C6 (Table S2) that met all criteria for being genuine T4SS inhibitor, but that, upon further evaluation, appeared to strongly affect *L. pneumophila* growth outside the host (Fig. 4A,B), suggesting that they likely had other molecular targets besides the T4SS. Inhibition of *L. pneumophila* growth in media did not require the presence of Dot/Icm system components as Lp02Δ26 remained sensitive to these two compounds (Fig. 4E). Interestingly, their growth-inhibitory activity was limited to *L. pneumophila* and did not affect other Gram-negative bacteria such as *E. coli* or *P. aeruginosa*, suggesting that these two compounds do not represent general antimicrobials but rather targeted an essential component or mechanism solely present in *L. pneumophila*.

Notably, none of the five hit compounds discovered here overlapped with the hits found in the study by Charpentier *et al.* (31). Contributing factors that may lead to this discrepancy in results likely include the different types of host cells used (RAW264.7 macrophages vs J774 macrophages), the aforementioned difference in the duration for which macrophages were exposed to the compounds prior to *L. pneumophila* challenge, as well as the criteria for selection of positive hits. Despite these differences, the T4SS inhibitors from either study set the stage for the development of a new generation of “smarter” antibiotics that, unlike most conventional antibiotics, could selectively target pathogens while leaving commensals unaffected. By exclusively altering the physiology of pathogenic bacteria that rely on a T4SS for virulence, such compounds have the potential to one day treat infectious diseases while preserving the healthy microbiota, a prerequisite for the prevention of secondary infections. Furthermore, if coupled to chemical handles, the compounds discovered here can be used as laboratory tools to further examine the regulation of bacterial T4SSs during infection. Although recent work has provided first insight into the structural organization of the *L. pneumophila* T4SS (14–18), much has yet to be learned about the dynamics of these sophisticated protein secretion machines.

## Acknowledgements

We thank all members of the Machner lab and Simeonov lab for their support and feedback. This work was supported by the Intramural Research Program of the National Institutes of Health, National Institute of Allergy and Infectious Disease (to RAH), National Center for Advancing Translational Sciences (to AS), and the *Eunice Kennedy Shriver* National Institute of Child Health and Human Development as well as the NIH Director’s Challenge Innovation Award (to MPM).

## Materials and Methods

### Strains, Media, and Reagents

*L. pneumophila* strains were grown and maintained as described (54). The bacterial strains used in this study are listed in Table S3. *L. pneumophila* strains Lp02 and Lp03 are thymidine-auxotroph derivatives of *L. pneumophila* strain Philadelphia-1 (55). *Legionella* were grown in ACES-buffered yeast extract with thymidine (AYET) supplemented at 100 μg/ml. Whenever indicated, chloramphenicol and kanamycin were used at a final concentration of 5 μg/mL and 20 μg/ml respectively. *E. coli* and *P. aeruginosa* were grown with aeration in Luria–Bertani (LB) medium. RAW264.7 macrophages were obtained from American Type Culture Collection and were grown in DMEM supplemented with 10% FBS and incubated in 5% CO_2_ at 37°C. β-lactamase-specific antibody was obtained from Abcam. Rabbit polyclonal anti-isocitrate dehydrogenase (ICDH) antibody was generously provided by Abraham (Linc) Sonenshein, Tufts University Medical School, Boston, MA.

### Plasmids

Plasmids and oligonucleotides used in this study are listed in Tables S4 and S5, respectively. Plasmids encoding β-lactamase effector protein fusions were constructed as previously described (35). The pXDC61 plasmid was generously provided by Howard A. Shuman (University of Chicago). Briefly, PCR products of the *lidA* flanking regions were digested with appropriate restriction enzymes and cloned into the KpnI-SmaI-BamHI-XbaI polylinker of pXDC61. Lp02 or Lp03 were transformed with plasmids by electroporation (56).

### Type IV translocation qHTS assay

Detection of β-lactamase-LidA fusion protein translocation into infected macrophages was performed using the FRET-based detection method described previously (35), with modifications. Briefly, Lp02 or Lp03 containing the plasmids that encode either β-lactamase or β-lactamase fusion proteins were grown to post-exponential phase in the presence of 0.5 mM IPTG. RAW264.7 macrophages were dispensed by a Multidrop Combi Reagent Dispenser (Thermo Scientific) into black clear-bottom 1536-well plates (Greiner Bio One) at 2×10^4^ cells/well and allowed to attach for 3 hours. The compounds assayed were solubilized in in DMSO and arrayed as five or six-point inter-plate titrations at final concentrations ranging from 46 μM to 0.18 μM, and 23 nL of compound solution was transferred into 1536 well plate using a Kalypsys pintool. After compound addition to macrophages, the cells were challenged with the appropriate strain of *L. pneumophila* at an MOI (multiplicity of infection) of 20 using a Multidrop Combi Reagent Dispenser. The CCF4/AM substrate (Life Technologies, Inc.) was added to the wells 1 h post-infection and then incubated for an additional 3 hours at room temperature. CCF4/AM fluorescence was measured using dual fluorescence intensities (Ex_1_=405±20, Em_1_=460±20, and Ex_2_=405±20, Em_2_=530±20 nm) in an EnVision Multimode Plate Reader (PerkinElmer, Boston, MA, USA). The ratio of fluorescence intensities (Em_1_/Em_2_) was calculated as representation for the β-lactamase activity levels within the host cells. When appropriate, individual assay wells were visualized using a Zeiss Observer.Z1 equipped with a βLac, blue/aqua fluorescence filter cube (Chroma Technology Corp).

### Compound libraries

The screening collection of 18,272 members included the following libraries with the number of compounds indicated in parentheses: NCATS Pharmaceutical Collection (NPC, 2816), NCATS Pharmacologically Active Chemical Toolbox (NIH, 2108), Mechanism Interrogation PlatE (NIH, 1912), Library of Pharmacologically Active Compounds ( LOPAC, 1280), Spectrum Collection (MicroSource Discovery Systems, 2000), Tocris (Tocris Bioscience, 1408), BioMol (Enzo Life Sciences, 1408), Cayman natural product (303), Pharmacopeia (3000), and KINACore Library (ChemBridge, 2037).

### β-lactamase specificity assay

Purified recombinant β-lactamase (ThermoFisher) in PBS (0.15 nM) was dispensed into 1536-well plates using a BioRAPTR FRD Microfluidic Workstation (Beckman Coulter Life Sciences). Compounds or control buffer (DMSO) were transferred using Kalypsys pintool equipped with a 1536-pin array. The plate was incubated for 10-15 min at room temperature, and nitrocefin (Calbiochem) was added to a final concentration of 430 μM by the BioRAPTR FRD workstation to initiate the color reaction. After a 20 min incubation at room temperature, hydrolysis of the nitrocefin substrate was measured by absorbance (OD_490_) using an Envision Multimode Plate Reader.

### Cytotoxicity assay

A Multidrop Combi Reagent Dispenser was used to dispense 5,000 or 50,000 RAW264.7 macrophages per well into either 1536- or 96-well opaque white plates (Greiner Bio One), respectively. The plates were incubated for 3 hours to allow for cell attachment to the substrate to occur. The compounds were transferred into 1536-well plates using a Kalypsys pintool or manually into 96-well plates. After 24 hours, Cell-Titer Glo (Promega) was added following the manufacturer’s instruction using a BioRAPTR FRD. The plates were incubated for 30 min at room temperature, and luminescence was measured using a ViewLux plate reader (Perkin Elmer).

### Intracellular growth assay

RAW264.7 macrophages were dispensed using a BioRAPTR FRD workstation at either 7,000 cells per well in 1536-well or 14,000 cells /well in 384-well plates with media supplemented with 0.5 mM IPTG. The cells were allowed to attach to the substrate for 3 hours before compounds were added using the Kalypsys pintool. The cells were immediately challenged at an MOI of 50 with Lp02 containing pXDC31 (GFP encoding plasmid) that grown to post-exponential phase in the presence of 0.5 mM IPTG. The plates were centrifuged at 200 x g for 5 min. After 14 hours incubation, the macrophages were chemically fixed for 15min at room temperature with formaldehyde (4% final concentration) and washed 2x with PBS. The cell nuclei were stained using Hoechst 33342 (Thermo-Fisher Scientific). Both 1536- and 384-well plates were imaged on an automated, widefield high content imager (IN Cell 2200, GE Healthcare) using a 10x/0.45 NA lens and standard DAPI (nuclear stain) and FITC (*Legionella* infection) excitation and emission filters. The TIFF files were quantitated using the canned Multi Target Analysis protocol (GE Investigator Workstation software v3.7.2). Briefly, nuclei from the DAPI channel were identified using top hat segmentation and a sensitivity setting of 96 and minimum size area of 35 μM^2^. Bacteria were detected in the FITC channel (cells) using a 2 μm collar dilation from the nuclear bitmap. FITC objects with an average nuclear RFU intensity above 450 (3Stdev above mean negative control wells) were considered as “Legionella positive”, and result were represented as relative infection rates [% of control].

### *In vitro* growth assay

Lp02, Lp03, JV4044 (a kind gift from Joseph Vogel, Washington University in St. Louis), *E. coli* K12 MG1655, and *P. aeruginosa* were diluted in the appropriate growth media to an OD_600_ between 0.01 to 0.1. The bacteria were treated with 28 μM of compound and loaded into 96-well plates at a total volume of 300ul per well. Growth was measured at an OD_600_ in one hour intervals for 18 hours or more in a Clariostar Monochromator Microplate Reader (BMG Labtech).

### Enumeration of Lamp1 positive LCVs when treated with compounds

RAW264.7 macrophages were grown in CellVis 8 chambered coverglass slides (#C8-1.5H-N) and challenged at an MOI of 25 with Lp02 producing mCherry from the plasmid pXDC50. DMSO or the indicated compounds (23 uM) were added concurrently with Lp02, slides were centrifuged at 200 x g for 5 min. After 2 hours of incubation, cells were washed 3x with PBS, fixed with formaldehyde (4% final concentration) for 10 min at room temperature and washed 2x with PBS. Staining for uninfected (outside) Lp02 was performed with rabbit-anti-*Legionella* primary antibody (1:3000, 1 hr, 37°C), cells were washed 3x with PBS, and stained with goat-anti-rabbit-404 secondary antibody (1:2000, 1 hr, 37°C, Life Technologies #C2764). Cells were then permeabilized with 0.2% TritonX 100 in PBS for 10 min at room temperature and washed with PBS. Cells were stained with rabbit-anti-Lamp1 primary antibody (1:1000, 1 hr, 37°C, Abcam #ab24170), washed 3x with PBS, and stained with goat-anti-rabbit-488 (1:2000, 1 hr, 37°C, ThermoFisher #656111). Slides were washed with PBS and internalized Lp02 with or without Lamp1 staining were quantified by microscopy using a Zeiss LSM 800 with Airyscan confocal microscope. Experiments were performed 3-4 times per compound, counting 50-100 internalized Lp02 each. Data were analyzed by dividing Lamp1 positive bacteria by the total number of internalized bacteria and represented as a percentage, averaging the percentages across replicates, and displaying the data as the mean with standard deviation and individual replicate points. A two-way unpaired T-test was performed comparing average counts of each compound to the DMSO control, (*) indicates P<0.05.

### Coxiella effector translocation assay and growth assays

*C. burnetii* CyaA translocation assays were performed using THP-1 macrophage-like cells (5 x 10^5^ per well) in 24-well plates as previously described (57). THP-1 cells treated overnight with 200 nM phorbol 12-myristate 13-acetate were washed once with growth medium (RPMI plus 10% FBS), infected with *C. burnetii* transformants expressing CyaA at an MOI ~ 50 for 24 h, and then incubated an additional 24h with fresh medium containing 50 μM of the indicated compounds. For CyaA translocation assays, the concentration of cAMP in lysates from infected THP-1 cells was determined using the cAMP enzyme immunoassay (GE Healthcare) as previously described (57). Positive secretion of CyaA fusion proteins by *C. burnetii* was scored as ≥2.5-fold more cytosolic cAMP than the negative control (wild-type *C. burnetii* producing CyaA alone) (57, 58).

In vitro replication of *C. burnetii* in the presence of compounds was determined as described (59). Briefly, 1 x 10^5^ per ml of *C. burnetii* were incubated at 37° C in a 2.5% O_2_ and 5% CO_2_ with either 5 μM of the indicated compounds or DMSO. The number of *C. burnetii* genomes measured by qRT-PCR in ACCM-D broth cultures was measured at the start of the experiment (Day 0) and after incubation (Day 6).

### Bacterial uptake assay

RAW264.7 macrophages (400,000/well) were seeded on coverslips in 24-well plates and grown overnight. The macrophages were treated with compound (57 μM final), Cytochalasin D (10 μM final), or DMSO for 5 min before the addition of Lp02 resuspended in tissue culture media containing compound (57 μM final), Cytochalasin D (10 μM final), or DMSO at an MOI of 1. The plate was centrifuged for 5 minutes at 200 x g to enhance bacteria-cell contact and incubated for 1h at 37°C in 5% CO_2_. Extracellular bacteria were removed by washing 3x with PBS. The cells were fixed in PBS containing 3.7% formaldehyde.

The cells were blocked in PBS containing 5% goat serum for 1 h at 37°C. Staining for uninfected (outside) Lp02 was performed with rabbit-anti-Legionella primary antibody, cells were washed 3x with PBS, and stained with goat-anti-rabbit FITC conjugated secondary antibody. Cells were then permeabilized with 100% methanol, blocked with 5% goat serum. Staining for total (inside and outside) Lp02 was performed with rabbit-anti-Legionella primary antibody, cells were washed 3x with PBS, and stained with goat-anti-rabbit Texas Red conjugated secondary antibody. After washing 3x with PBS, the cell nuclei were stained using Hoechst 33342, and the coverslips were mounted on slides before analysis.

## Supplemental Figures and Tables

**Supplemental Figure 1:**

The production of βLac or βLac_LidA was induced overnight by incubation with IPTG in both strains Lp02 and Lp03. Proteins within the bacterial lysate were separated by SDS-PAGE. After transfer to nitrocellulose membranes, the βLac-tagged proteins were immunologically detected using a β-lactamase specific antibody (top). Isocitrate dehydrogenase (ICDH) served as a loading control (bottom).

**Supplemental Figure 2:**

(A) β-lactamase specificity assay principle based on hydrolysis of the chromogenic substrate nitrocefin, an indicator of β-Lac enzymatic activity. (B) Cell cytotoxicity assay principle based on ATP abundance, an indicator of metabolically active cells.

**Supplemental Figure 3:**

Legionella growth inhibition in a 1536 well plate format. RAW264.7 macrophages were challenged with a GFP-producing *L. pneumophila* for 14 hours in the presence of the indicated compounds, and the GFP signal (as a measure for bacterial growth) was determined using an IN Cell Analyzer 2200 Imaging system.

**Supplemental Figure 4:**

Bacterial uptake by macrophages in the presence of compounds. RAW264.7 macrophages were treated with the indicated compounds, Cytochalasin D, or DMSO vehicle prior to infection with Lp02, and intracellular bacteria were enumerated visually. Data shown are the mean with standard deviation from six experimental replicates, with the DMSO control considered as 100%. P-values (t-test) are <0.05 (*), <0.001 (**), or <0.005 (***) relative to the control.

**Supplemental Figure 5:**

In vitro replication of *C. burnetii* in the presence of compounds. ACCM-D broth cultures containing 5 μM of the indicated compounds or DMSO and inoculated with ~ 1 x 10^5^ per ml of *C. burnetii* were incubated at 37° C in a 2.5% O_2_ and 5% CO_2_. The plot depicts the number of *C. burnetii* genomes measured in ACCM-D broth cultures at the start of the experiment (Day 0) and after incubation (Day 6) detected by qRT-PCR *(1)*. Bars indicate the mean ± SEM of combined data from three independent experiments each performed with triplicate samples (N = 9). One-way ANOVA analysis indicated differences between groups at D6 were found not significant (ns) using a 95% confidence interval.

**Table S1 - Data of 113 compounds selected from the qHTS reporter translocation assay. \*** Asterisk indicates compounds that have isomers

**Table S2 – Data of 7 compounds selected following validation assays.**

**Table S3 – Strains used in this study.**

**Table S4 – Plasmids used in this study.**

**Table S5 – Oligonucleotides used in this study.** Underlined sequences represent restriction enzyme sites.

